# When exploration replaces storage: how eye movements shape visual working memory

**DOI:** 10.64898/2026.01.04.697560

**Authors:** Rania Qais, Robert T. Knight, Shlomit Yuval-Greenberg

## Abstract

Visual working memory (VWM) is traditionally studied while constraining eye movements and limiting access to visual input, yet in natural vision humans constantly explore and resample their environment. Only a few studies have examined VWM utilization when participants were allowed to interact with the environment and found that participants often preferred to resample their environment rather than rely on VWM storage. However, since eye movements were not controlled in these studies, the link between VWM utilization and free visual exploration remained unknown. In two experiments (N = 40), we investigated how visual exploration shapes reliance on VWM versus perceptual input. Participants searched for a common target across two item sets and could either store multiple items for comparison or repeatedly resample the sets by switching between them. Results revealed that when switching was achieved through eye movements, participants consistently relied more on visual resampling and less on VWM; in contrast, when switching required a manual response, they shifted toward greater VWM use. This pattern persisted even when peripheral input was equated, suggesting that natural exploration through eye movements reduces the cognitive cost of acquiring visual information, leading to a strategic reduction in VWM use. Our findings challenge fixation-based approaches to VWM research and highlight the importance of studying cognition under ecological viewing conditions.

## Introduction

Visual working memory (VWM) is the cognitive system that stores and manipulates visual information over short intervals (Luck & Vogel, 1997). Over the past few decades, research has thoroughly investigated VWM capacity, asking how much visual information can be stored and kept accessible for ongoing tasks (Luck & Vogel, 2013). Although the precise capacity of VWM is still under debate and depends on several factors (Cowan, 2001), including item type (Luria & Vogel, 2011a) and individual ability (Luria & Vogel, 2011b; Vogel et al., 2005)(Cowan, 2001), evidence suggests that the typical capacity in healthy adults is around 3-4 items in most cases (Chawoush et al., 2023; Sperling, 1960; Vogel et al., 2001).

However, despite decades of research, relatively little is known about how VWM is used in everyday situations in which observers are free to move and interact with their environment (Draschkow et al., 2021). This gap stems from the fact that most VWM studies are highly controlled and focus on situations where perceptual input disappears and reappears while participants’ information-seeking behaviors are restricted. For example, in the classic change detection task, participants are asked to maintain fixation while items appear briefly, then disappear for a short delay, and later must be retrieved from VWM for comparison or manipulation (Luck & Vogel, 1997). This approach is well-suited for assessing maximal VWM capacity, as it pushes the system to its limits. It is, however, unnatural for two reasons: (a) in natural life, stimuli do not simply appear out of nowhere and then vanish - they typically appear because the eyes moved and landed on them and disappear because the eyes moved away from them; and (b) humans are inherently exploratory, frequently interacting with their environment and rarely keeping their eyes still for more than a few hundred milliseconds (Yarbus, 1967). Whether these constraints limit the generalizability of standard VWM findings outside the laboratory remains an open question.

Only a handful of studies have examined VWM in more naturalistic contexts, where participants can actively access and resample visual information (Ballard et al., 1995; Draschkow et al., 2021; Kvitelashvili & Kessler, 2024; Li et al., 2023; Sahakian et al., 2023, 2025; Somai et al., 2020). These studies consistently show that when visual information is readily available, participants tend to rely more on perceptual resampling of external input than on internal storage, preferring to resample repeatedly rather than use stored representations and thereby underutilizing VWM capacity. Yet, despite the growing literature on this topic, the mechanisms governing VWM use in everyday viewing remain unknown. In particular, since none of these previous studies examined the isolated effects of eye movements on VWM, it remains unknown what role free visual exploration plays in this process. Given that visual exploration via eye movements is one of the most fundamental human behaviors, and that almost everything we know about VWM is based on tasks that limit eye movements, examining the link between VWM and eye movements in free exploration is essential for understanding this function.

Here, we test the hypothesis that visual exploration driven by eye movements plays a critical role in VWM utilization. We propose that the relation between the availability of visual information and reliance on VWM depends on natural viewing: when input can be accessed through eye movements, observers prefer to sample the environment directly and rely less on VWM; but, when access is unnatural and non-exploratory, they instead rely more heavily on VWM. This hypothesis was examined using a task inspired by the game Dobble (Spot It! in the US).

In two experiments, participants were asked to identify a common item between two sets, presented either simultaneously, allowing free eye movements between them, or sequentially in the center with switching via button presses. They could adopt one of two strategies: (1) store multiple items from one set into VWM and then compare them with the other set, or (2) switch frequently between sets to refresh perceptual input and reduce reliance on memory. The findings revealed underutilization of VWM only when switching was achieved by eye movements, a natural exploratory action, but not when switching required button-presses. These findings indicate that VWM use under free-viewing differs from that measured under constrained fixation and underscore the value of studying cognition in unconstrained, ecologically valid settings.

## Experiment 1

### 1. Methods

#### 1.1. Participants

Twenty participants (16 females; age range 20-34, M = 24.67, SD = 3.29) took part in Experiment 1. The sample size was determined based on a power analysis, using G*Power 3.1. (Faul et al., 2007) . We examined the main effect of display condition on the gathering variable as a dependent-samples two-tailed t-test, with α = .05, targeting 90% power to detect a main effect of Cohen’s d = 1. This led to a minimal requirement of 13 participants, which we expanded to 20. The experiment was approved by the University’s Ethics Committee, and all participants signed an informed consent form. Participation was in exchange for either course credit or payment of the equivalent to approximately 10 USD. All participants reported normal or corrected-to-normal vision and no history of neurological disorders.

#### 1.2. Stimuli

The stimulus set consisted of 184 colored drawings of familiar visual objects (size 2.6°-2.7°), chosen from Snodgrass and Vanderwart’s object pictorial set (Snodgrass & Vanderwart, 1980) and Multipic database (Duñabeitia et al., 2018). In each trial, such 3-7 objects were presented on large white circles referred to as “cards” (diameter 9°). Objects were spatially distributed across the card surface. The background was always mid-gray.

#### 1.3. Procedure

Participants were seated in a sound-proof chamber, with their head resting on a chinrest at a distance of 100 cm from the display monitor (24” LCD ASUS VG248QE, 1,920 × 1,080 pixels resolution, 120 Hz refresh rate, mid-gray luminance 110 sd/m2). The experiment was generated and controlled using PsychoPy software in Python. The procedure and design parameters were examined and validated in a preliminary study.

Each trial began with a central black fixation cross (0.8° × 0.8°) presented for 1.5 seconds, which participants were asked to fixate. This was followed either by the presentation of two cards placed symmetrically to the left and right of center at a distance of 7.72° from each other (simultaneous condition) or by the presentation of one central card, which could be switched to another centrally presented card by pressing either the right or the left key (sequential condition). The distance between the cards in the simultaneous condition was determined based on a preliminary study.

In the sequential condition, each key press was followed by a 50 ms blank screen, after which the second card was presented. Pressing the key again, resulted in another 50 ms blank screen, followed by the first card, and so on. This brief blank is included to equate the sequential condition with the simultaneous condition, in which a saccade of similar duration occurs. Since saccades involve saccadic suppression and blurred visual input, the blank screen mimicked this perceptual gap.

In both conditions, the pair of cards (presented either simultaneously or sequentially) contained the same set size (3, 5, or 7) and exactly one common item (the target). Participants could move their eyes freely to search for the common item and were instructed to press the spacebar as soon as they identified it. After pressing the spacebar, the two cards disappeared and a screen displaying randomly selected items from that trial (including the target) appeared, with each item numbered. Participants were instructed to press the corresponding number key to indicate the target. The two conditions were presented in separate blocks, with order counterbalanced between participants. The main experimental session consisted of 10 blocks of 45 trials each - 5 blocks for the simultaneous condition and 5 blocks for the sequential condition. The trial procedure of both conditions is illustrated in Fig. 1.

**Figure 1.**
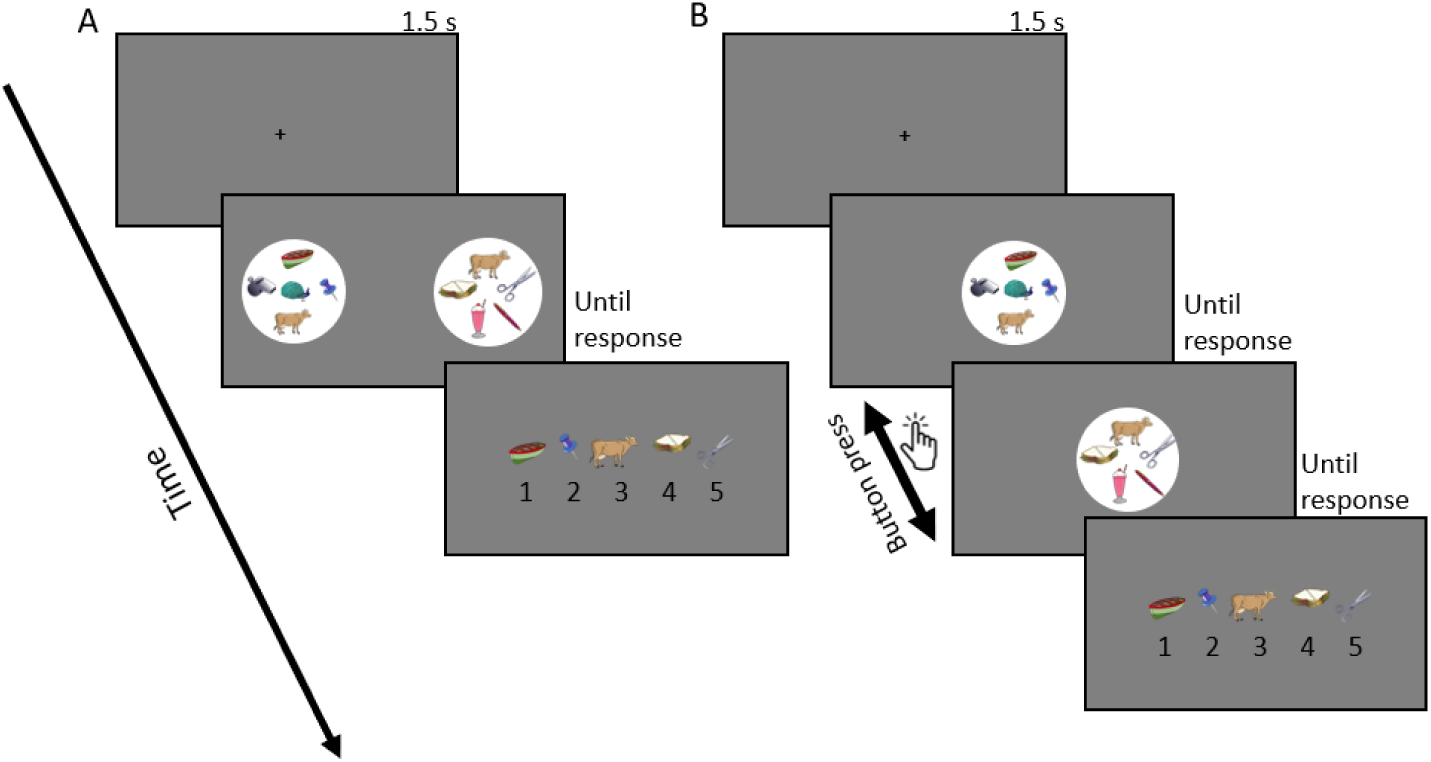
Procedure of Experiments 1. **(A)** Simultaneous Condition. Each trial began with a black fixation cross, followed by a presentation of two cards (white circles), with 3,5, or 7 colored items, and distances of 7.72° between the cards. Participants were free to move their eyes to search for the target. Once the target was identified and the spacebar was pressed, the screen advanced to a display showing items from that trial, arranged horizontally on the screen with numbers underneath. The correct answer in this case is 3 representing the object that was the common item between the two cards **(B)** Sequential Condition: As in the simultaneous condition, but only one card was presented in the center of the screen, with 3,5, or 7 colored items. Participants were free to move their eyes within the presented card and could switch between the two cards by pressing a key. Once the target was identified and the spacebar was pressed, the screen advanced to the response screen as in the simultaneous condition and participants were asked to press the number representing the target item (3 in this case). Figures are not to scale for clarity of presentation

#### 1.4. Eye tracking

Binocular gaze position was monitored using a remote infrared video-oculographic system (Eyelink 1000 Plus; SR Research, Canada), with a spatial resolution <0.01° and average accuracy of 0.25°-0.5° using a chinrest (manufacturer specifications). Raw gaze positions were sampled at 1000 Hz and converted into degrees of visual angle using a 9-point-grid calibration, performed at the start of the session and repeated between blocks when necessary.

#### 1.5. Data analysis

##### 1.5.1. Behavioral analysis

Mean reaction times (RT) and accuracy rates were calculated separately for each participant and condition. RT - defined as the time from card presentation to spacebar press - was calculated only for correct trials. Accuracy rates represented the proportion of correct target identifications in each condition.

##### 1.5.2. Eye movement analysis

Gaze positions were segmented from trial onset and until the spacebar press. Eye tracking samples with lost pupil tracking, due to blinks or faulty recording, were interpolated. Saccades were detected using a published algorithm (Engbert & Kliegl, 2003). Only saccades exceeding 1° of visual angle were included, in order to exclude microsaccades within single items and focus on gaze shifts between items. Eye tracking analysis focused on two main measurements: the switching-rate index, reflecting how often participants switched between the two cards, and the gathering index, reflecting how many items from one card were gathered into working memory before switching to the next card. The two conditions differed in the mean by which switching was done: while in the sequential condition each button press counted as a switch, in the simultaneous condition, a switch was defined as a large saccade crossing the vertical midline of the screen. In both cases, switching-rate was calculated per trial as the number of switches divided by trial duration in seconds. The gathering index was calculated similarly for the two conditions and defined as the average number of saccades performed within one card (i.e. did not cross the vertical midline) between two switches, including saccades from trial onset until the first switch. Both indices were calculated per trial and then averaged per participant and condition. Example eye movement patterns are shown in Fig. 2.

**Figure 2.**
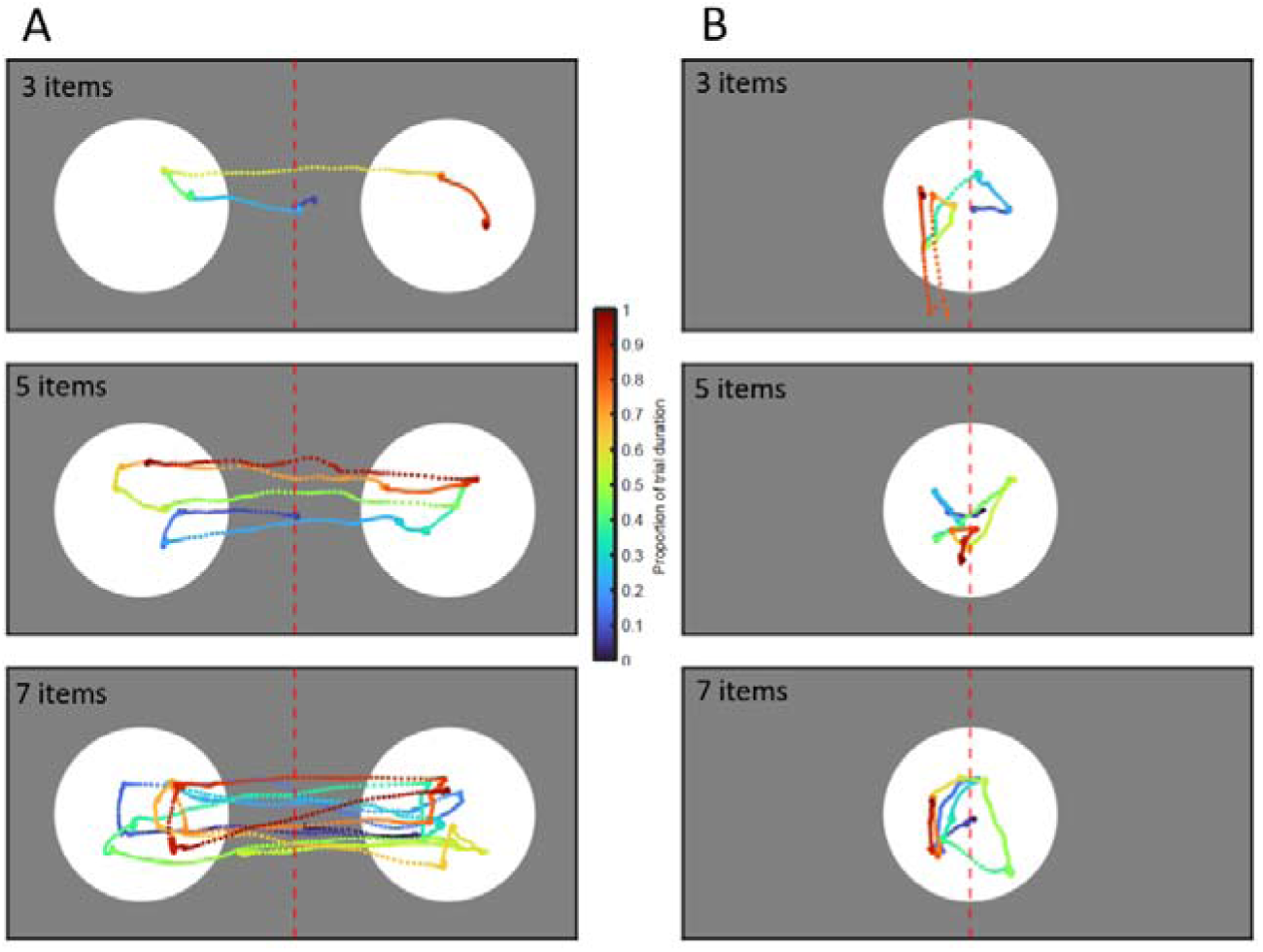
Example gaze traces of Experiments 1. Gaze trace of six example trials in Experiment 1 with three items (top), five items (middle), and seven items (bottom), in the simultaneous (A) and sequential (B) conditions. The gray rectangle represents the screen, with the white circles representing the cards. The red dotted line represents the midline and was not presented to the participants. Saccades that crossed the midline were considered switches in the simultaneous condition, and saccades within a circle were considered as reflecting gathering. The colors represent trial progression, with blue representing gaze trace early during the trial, green representing mid-trial and dark red representing the end of the trial.

##### 1.5.3. Analysis software

Eye movements and behavioral analysis were processed in MATLAB R2022b (The MathWorks Inc., Natick, MA, USA). Statistical analysis was done in JASP 0.19.1.

##### 1.5.4. Data availability

The datasets generated and analyzed during the current studies, along with the MATLAB script used for analysis and visualization, are available in the Open Science Framework (OSF) repository at: https://osf.io/w547b/overview

## 2. Results

Accuracy rates, RTs, gathering index, and switching-rate index were analyzed separately using four Repeated Measures 3X2 ANOVA, with set size (3, 5 or 7) and display condition (simultaneous or sequential) as factors.

### 2.1. Accuracy rates

Accuracy rates were high (grand average ranging between 0.965 to 0.991), limiting the ability to interpret differences. Nevertheless, the analysis revealed a main effect of set size (F(2,38) = 7.062, p = .002): when the set size increased, accuracy rates decreased (negative linear trend: t(19) = -3.150, p = .005). A significant main effect for display condition was also found (F(1,19) = 6.226, p = .022), indicating that participants were less accurate in finding the right target in the sequential than in the simultaneous condition (t(19) = -3.143, p = .005). There was an interaction between the two factors, suggesting that the effect of set size on accuracy rates differed between the display conditions (F(2,38) = 4.267, p = .021). While there was a significant effect in the sequential condition (main effect F(2,38) = 6.731, p = .003; negative linear trend: t(19) = -3.143, p = .005), no such effect was found in the simultaneous condition (main effect F(2,38) = 1.130, p = .334; Fig. 3A).

**Figure 3.**
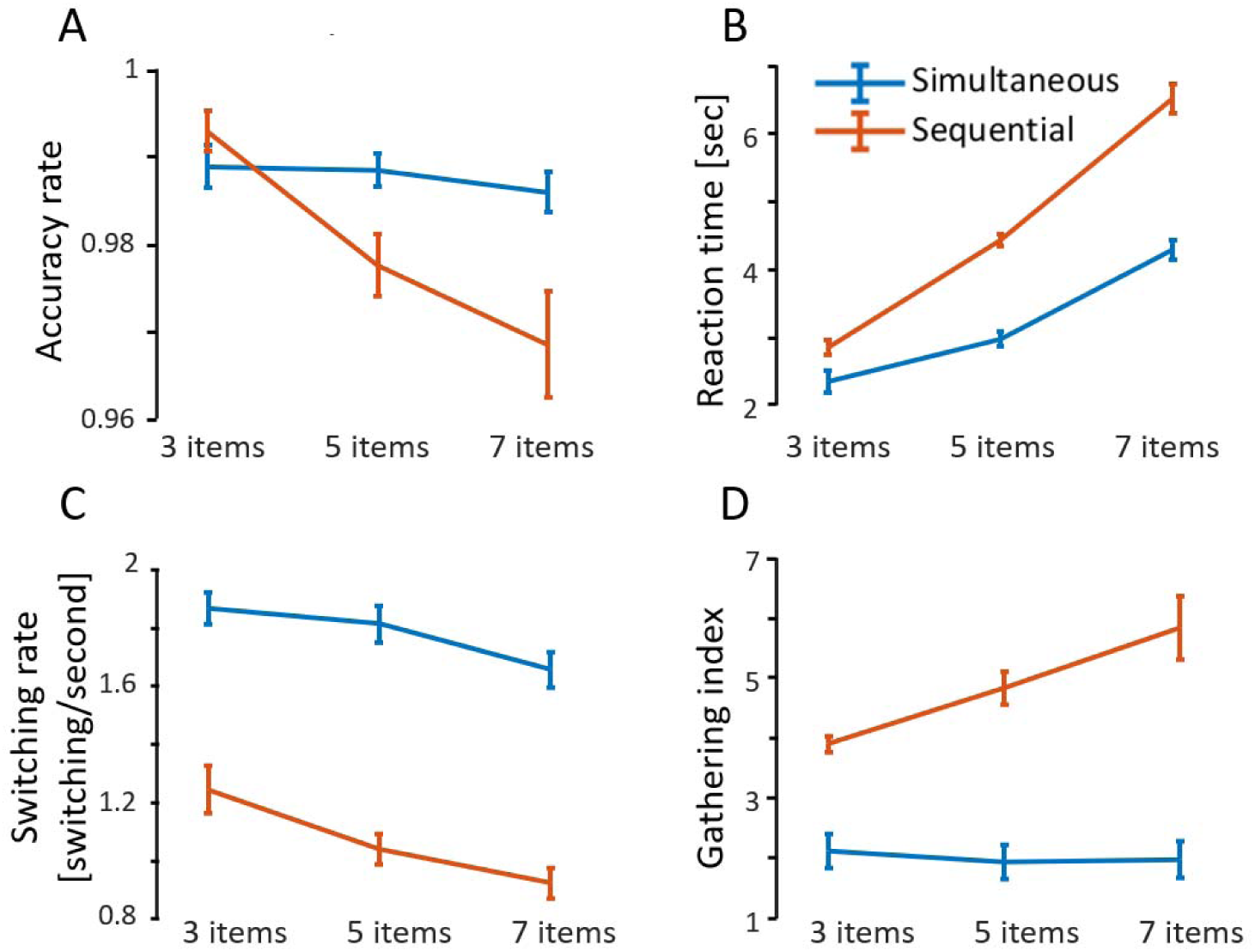
Results of Experiment 1. **(A)** Accuracy-rates averaged across participants, within each condition. Accuracy rates were overall higher in the simultaneous than in the sequential condition. As the set size increased, accuracy in the sequential condition decreased, whereas accuracy in the simultaneous condition remained unchanged. **(B)** Mean reaction times (RTs) were calculated on correct trials and averaged across participants, within each condition. RTs were faster in the simultaneous than in the sequential condition and increased with set size in both conditions. **(C)** Switching-rate is expressed as the mean number of switches per second, averaged across participants within each condition. Switching rate was higher in the simultaneous than in the sequential condition and decreased with set size in both conditions, with a steeper trend in the sequential condition. **(D)** Gathering index is expressed as the average number of small saccades performed on one set of items before moving to the next. Gathering was higher in the sequential than in the simultaneous condition and increased with set size in both conditions. Error bars in all panels represent the within-subject standard error Cousineau, 2005).

### 2.2. Reaction times

As expected, larger set sizes resulted in longer RTs (main effect of set size: F(2,38) = 293.340, p < .001; positive linear trend: t(19) = 17.962, p < .001). Participants took longer to find the target in the sequential condition than in the simultaneous condition (main effect of display condition: F(1,19) = 59.688, p < .001). A significant interaction indicated that the effect of set size on RT varied between display conditions (F(2,38) = 11.739, p < .001). In both conditions there was a significant main effect of set size (simultaneous: F(2,38)= 232.388, p < .001; sequential: F(2,38) = 152.495, p < .001) with a positive linear trend (simultaneous: t(19)= 17.512, p < .001; sequential: t(19) = 12.833, p < .001), but the trend was steeper in the sequential condition. These findings indicate RT and accuracy effects are consistent with each other and are not due to speed-accuracy trade-offs. See Fig. 3B.

### 2.3. Switching-rate

The analysis revealed a main effect of set size (F(2,38) = 97.227, p < .001), indicating that as set size increased, participants performed fewer switches per second, either saccades in the simultaneous condition or key presses in the sequential condition (negative linear trend: t(19) = -12.626, p < .001). One-way ANOVAs performed separately for each display condition revealed a significant effect of set size in both (simultaneous: F(2,38) = 79.410, p < .001; sequential: F(2,38) = 48.054, p < .001). There was also a main effect of the display condition (F(1,19) = 37.042, p < .001), with a higher rate of switching in the simultaneous condition than in the sequential condition. There was no evidence for an interaction between the factors (F(2,38) = 1.985, p = .151; Fig. 3C).

### 2.4. Gathering

The simultaneous condition showed gathering values of 1.3-1.6, which are lower than the known VWM capacity of 3-4 items (Chawoush et al., 2023; Sperling, 1960; Vogel et al., 2001). This suggests that VWM capacity was not fully utilized in this condition. In contrast, in the sequential condition, where cards were switched by button presses rather than eye movements, gathering values were much higher (3.56, 4.64, and 5.57 for 3, 5, and 7 items, respectively), and closer to the typical VWM capacity, suggesting increased reliance on VWM. Consistent with this observation, there was a significant main effect of display condition, with higher gathering index for the sequential condition (F(1,19) = 27.004, p < .001; Fig. 3D). Despite a significant interaction between set size and display condition (F(2, 38)= 11.647, p < .001), this effect was found for set sizes when analyzed separately (3 items; F(1,19) = 33.078, p < .001; 5 items: F(1,19) = 28.607, p < .001; 7 items (F(1,19) = 22.136, p < .001).

We also found a significant main effect for set size (F(2,38)= 18.888, p < .001), with gathering positively associated with set size in both display conditions (simultaneous: F(2,38)= 48.147, p < .001, positive linear trend - t(19) = 7.606, p < .001; sequential: F(2,38) = 15.161, p < .001, positive linear trend - t(19) = 4.038, p < .001). However, this trend was steeper in the sequential tha in the simultaneous condition, suggesting that reliance was more affected by load in the sequential than in the simultaneous condition.

### 2.5. Summary of Experiment 1 results

Experiment 1 compared two display conditions: the simultaneous condition, with two cards presented side-by-side, and the sequential condition, with only one card presented at a time and switching taking place via a button-press. The simultaneous condition mimics natural viewing, in which eye movements can be used to collect perceptual data and compare it. In contrast, although there was no fixation requirement in the sequential condition, switching could not be done through eye movements, making it similar to a controlled lab setting in which participants keep their eyes stable while stimuli appear and disappear.

The findings show that in the sequential condition, participants performed more gathering and less switching than in the simultaneous condition. This indicates that in the simultaneous condition, which is closer to natural viewing, there is less reliance on VWM and more on foveal perceptual input. In other words, participants preferred to switch their gaze back and forth between the cards to compare the items one by one, rather than fixate at a few items within one card, store them in VWM, and then compare them to items on the other card.

We also find that set size affected the two display conditions similarly: with more items per card, participants tended to gather more and switch less. However, in the sequential condition, the gathering trend was stronger than in the simultaneous condition. In the simultaneous condition, the change in gathering between 3, 5, and 7 items is minimal. Crucially, even with a relatively large set of 7 items, participants preferred to shift their gaze between the cards rather than store items for later comparison.

Findings of Experiment 1 supported our hypothesis that, in the context of natural viewing, observers prefer to rely on perceptual input rather than on VWM storage. However, there was one crucial difference in perceptual input between the two display conditions: in the sequential condition, participants viewed only one card at a time, whereas in the simultaneous condition, they viewed both. This means that in the simultaneous condition, peripheral perceptual input was available, whereas in the sequential condition it was not. An alternative explanation for the findings is that participants prefer to switch more and gather less when peripheral information is available.

The second experiment addressed this question by providing identical peripheral input in both conditions. In the sequential condition of this experiment, two cards were presented: one at the center and one in the periphery. Participants were instructed to fixate only on the central card, and this was enforced with a gaze-contingent procedure. They could switch between the cards using the left and right arrow keys. Upon pressing the key, the central card moved to the periphery, and the peripheral card moved to the center (keeping the right card always on the right and the left on the left). This manipulation mimicked the perceptual change caused by a saccade in the simultaneous condition, while preserving an equivalent para-foveal and peripheral input.

## Experiment 2

### 3. Methods

The method of Experiment 2 was identical to that of Experiment 1, except where indicated otherwise.

#### 3.1. Participants

Twenty participants (13 females; age range 19-25, M = 22.2, SD = 1.63) took part in Experiment 2.

#### 3.2. Stimuli

The stimuli were the same as in Experiment 1, except that the cards and the objects were smaller (card diameter 4.58°; object size: 0.98°) and the objects were in grayscale.

#### 3.3. Procedure

After Experiment 1, we replaced the display monitor with a 24.5” BENQ XL2566K (1920×1080 pixels resolution, 360 Hz refresh rate). This experiment was almost identical to Experiment 1, except that the two cards were presented simultaneously even in the “sequential” condition. In the sequential condition, one card was presented at the center of the screen, and the other was placed 7.72° to the right or left of fixation. Participants were asked to look only at the central card and received a real-time warning if they looked away. Upon a key press, the two cards changed positions in the following way: the peripheral card shifted to the center and the central card shifted to the opposite peripheral side to preserve the distance of 7.72° as shown in Fig. 4. This design change was intended to mimic the retinal image shift caused by a saccade in the simultaneous condition, thereby controlling for the effects of peripheral vision in the two conditions.

**Figure 4.**
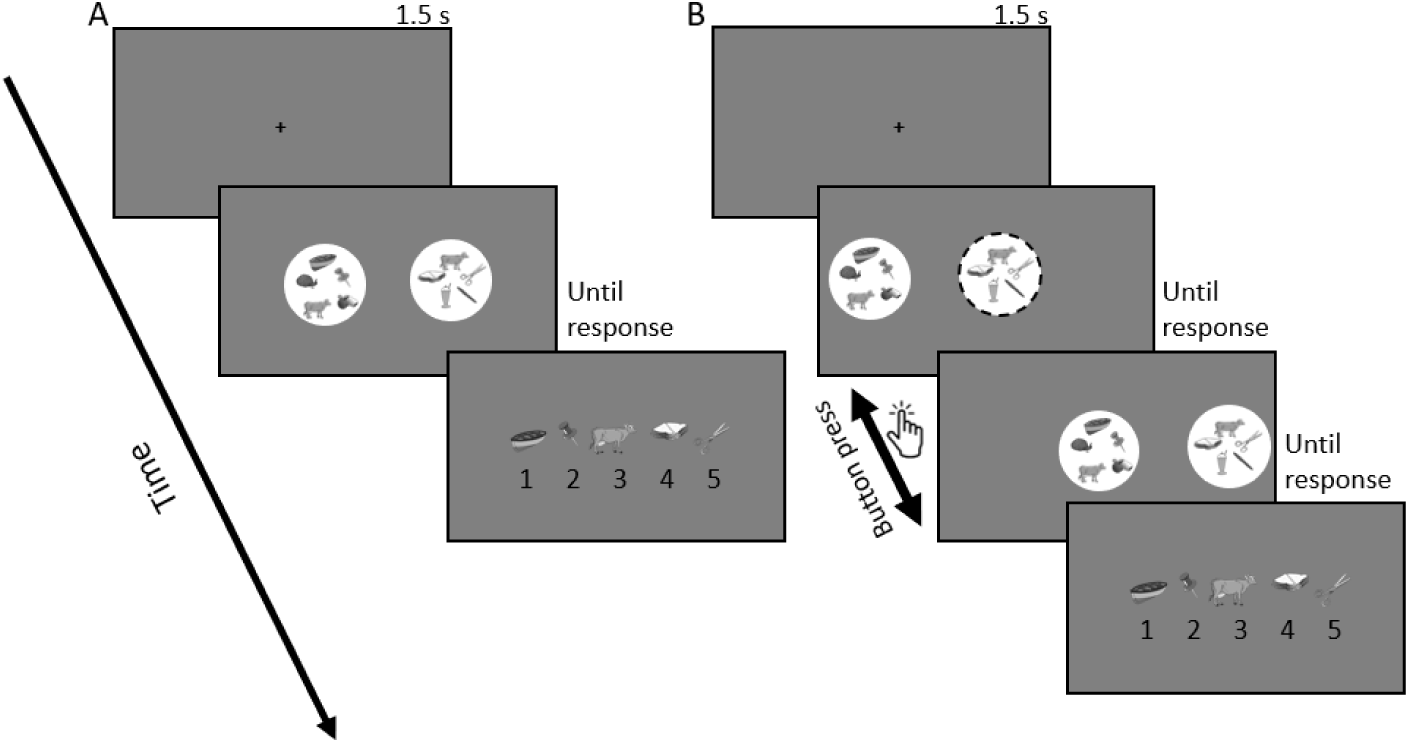
Procedure of Experiment 2. **(A)** Simultaneous Condition: Participants viewed two cards with 3, 5, or 7 items (black and white) and were free to move their eyes to search for the target. In this case, gaze switched from the right card to the left. After identifying the target and pressing the spacebar, items from the trial were displayed horizontally with identifying numbers. **(B)** Sequential Condition: One card appeared centrally and the other peripherally (left/right). Participants initially viewed only the central card. After a key press, the cards shifted sides to maintain a constant distance between them. Participants could switch between the two cards using a key. Figures are not to scale for clarity of presentation.

### 4. Results

#### 4.1. Accuracyrates

Due to an error in the experiment code, response correctness was not saved for 12 out of 20 participants. For the 8 participants with available accuracy data, accuracy was high in all conditions, as in the first experiment (range: 0.957-0.991; Fig. 5A). The analysis of these participants produced results quantitatively identical to those of Experiment 1. Accuracy was negatively associated with set size (main effect: F(2,14) = 4.877, p = .025; negative linear trend: t(7) = -2.805, p = .026), but this effect was modulated by display condition (Interaction: F(2,14) = 4.407, p = .033) and was found significant only in the sequential condition (main effect: F(2,14) = 5.114, p = .022; negative linear trend: t(7) = -2.996, p = .020) and not in the simultaneous condition (F2,14) = 2.404, p = .127). As in Experiment 1, accuracy was higher in the simultaneous than in the sequential condition (main effect: F(1,7) = 14.537, p = .007; Fig. 5A).

**Figure 5.**
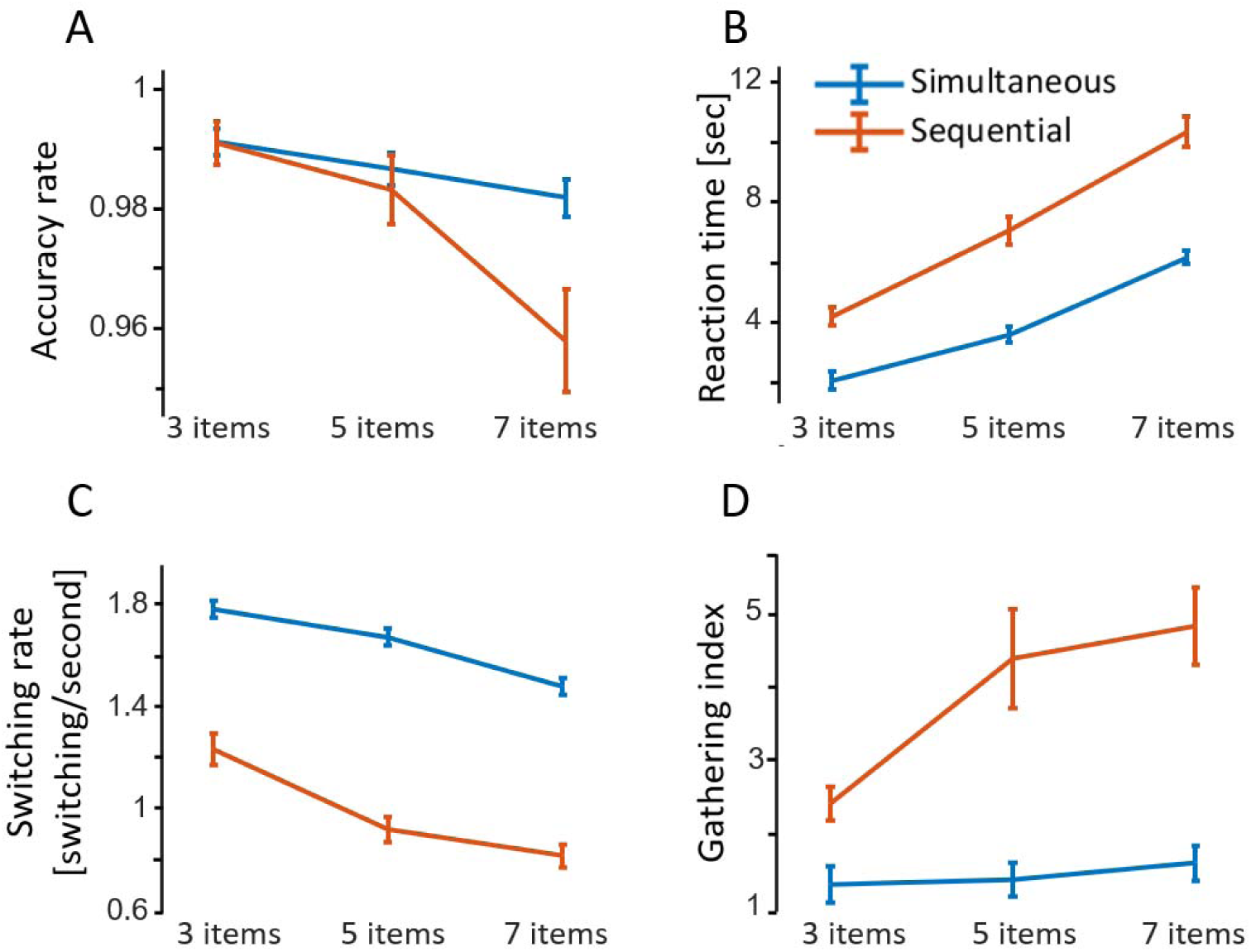
Results of Experiment 2. (A) Accuracy rates averaged across 8 participants who had accuracy data. Accuracy was overall higher in the simultaneous than in the sequential condition. As set size increased, accuracy in the sequential condition decreased, whereas accuracy in the simultaneous condition remained unchanged. **(B)** Mean RTs averaged across all trials for all participants. RTs were faster in the simultaneous than in the sequential condition, and increased with set size, in both display conditions. **(C)** Switching rate is expressed as the mean number of switches per second, averaged across participants, within each condition. Switching rate was higher in the simultaneous than in the sequential condition and decreased with set size in both conditions, with a steeper trend in the sequential condition. **(D)** Gathering index is expressed as the average number of small saccades performed on one set of items before moving to the next. Gathering was higher in the sequential than in the simultaneous condition and increased with set size in both conditions, with a steeper trend in the sequential condition. Error bars in all panels represent the within-subject standard error (Cousineau, 2005).

#### 4.2. Reaction times

In Experiment 1, RTs were calculated only for correct trials. However, since this information was missing for 12 participants and since accuracy in the rest of the participants was high, RTs were analyzed for all trials, assuming that most were correct. The results were qualitatively identical to those of Experiment 1. There was a significant main effect for the set size (F(2,38) = 108.970, p < .001) described by a positive linear trend (t(19) = 12.485, p < .001). There was also a significant main effect of display condition (F(1,19) = 51.060, p < .001), indicating that RT was longer in the sequential compared to the simultaneous condition. A significant interaction (F(2,38)= 4.635, p = .016) indicated that the effect of set size on RT was modulated by the display condition: set size had a positive effect on RTs in both conditions (simultaneous: F(2,38)= 130.732, p < .001, positive linear trend: t(19)= 12.470, p < .001; sequential: F(2,38) = 46.058, p < .001; positive linear trend: t(19) = 8.810, p < .001), but the effect was steeper in sequential condition (Fig. 5B).

#### 4.3. Switching-rate

As in Experiment 1, there was a main effect for set size (F(2,38)= 37.023, p < .001), indicating a lower switching rate (saccades or key presses) as set size increased. This negative association was found in both display conditions (simultaneous: F(2,38) = 134.276, p < .001 negative liner trend: t(19) = -13.790, p < .001; sequential: F(2,38) = 17.233, p < .001, negative linear trend: t(19) = -4.833, p < .001), but a significant interaction (F(2,38)= 4.577, p = .017), revealed that it was steeper in the sequential condition. There was also a main effect of the display condition (F(1,19) = 109.451, p < .001), with more switches in the simultaneous condition than in the sequential condition. This effect was found to be significant for all set sizes (3 items: F(1,19) = 48.499, p < .001; 5 items: F(1,19) = 96.660, p < .001, and 7 items: F(1,19) = 102.845, p < .001; Fig. 5C).

#### 4.4. Gathering

As in Experiment 1, there was a main effect of set size (F(2,38)= 6.302, p = .004), modulated by the display condition (interaction: F(2, 38)= 4.473, p = .018). When analyzed separately, set size had a positive linear effect in both conditions, but this effect was steep in the sequential condition and minimal in the simultaneous condition (simultaneous condition: main effect: F(2,38)= 64.595, p < .001, positive linear trend: t(19) = 9.096, p < .001; sequential: main effect: F(2,38) = 5.332, p = .009, positive linear trend: t(19) = 3.728, p = .001). A significant main effect for the display condition (F(1,19) = 26.212, p < .001) revealed that participants performed more gathering in the sequential than in the simultaneous condition, suggesting that they relied more on VWM in that condition. This was true for all set sizes (3 items: F(1,19) = 24.708, p < .001; 5 items: F(1,19) = 11.946, p = .003; 7 items: F(1,19) = 21.251, p < .001). See Fig. 5D.

#### 4.5. Summary

The results of Experiment 2 replicated those of Experiment 1 across all four measurements, confirming that the findings cannot be explained by peripheral input.

## 5. Discussion

The present study examined how exploration using eye movements shapes reliance on VWM. Findings showed that the choice of strategy depended on the type of motor action: when switching was done via eye movements, participants relied minimally on VWM, typically storing only 1-2 items before making a saccade to the other set; in contrast, when switching required a button press, participants tended to store more items (around 3-5) before switching, indicating greater reliance on VWM.

### 5.1. The effect of eye movements on VWM utilization

Previous research has shown that when visual information is easy to access, observers prefer to resample the environment rather than rely on stored presentations (Ballard et al., 1995; Draschkow et al., 2021; Droll & Hayhoe, 2007; Kvitelashvili & Kessler, 2024; Li et al., 2023). Our findings align with this conclusion but refine it by showing that reliance on VWM in natural contexts depends on the type of motor action used to obtain visual information: when participants used eye movements to obtain visual input, they tended to rely on perceptual resampling rather than VWM storage; when they were required to use button presses to access the same information, they relied more heavily on VWM. This pattern was evident even when we fully controlled for peripheral input across conditions, so the only difference was the means of interaction - eye movement versus button press. This indicates that neither the presence of perceptual input nor the ability to access it actively is sufficient to render VWM redundant; redundancy arises when available input can be accessed naturally through exploratory behavior such as eye movements. Thus, eye movements have a unique effect on VWM utilization: VWM is underutilized when visual input is accessible, but only when access occurs via eye movements.

The interplay between the cost of accessing visual information and the benefit it provides is a critical factor in determining reliance on VWM. When access is easy, participants prefer resampling over memory storage (Draschkow et al., 2021; Somai et al., 2020), and as access becomes more effortful, they increasingly rely on VWM. Sahakian et al. (Sahakian et al., 2023) suggested that this cost-benefit balance reflects not only changes in encoding but also changes in the decision threshold for using stored information rather than resampling the environment. Eye movements (and perhaps also reaching movements, as in Draschkow et al., 2021) differ from button presses in their attentional cost and in how they influence reliance on VWM. Although button presses are not goal-directed in space and, therefore, involve simpler preparatory calculations, they likely impose greater attentional demands in the context of a search task. Arguably, the difference in the attentional demands of these two behaviors reflects the fact that eye movements are automatic, hard-wired behaviors used constantly to explore the environment, whereas pressing a button to reveal input is a learned behavior that is novel in evolutionary terms. When weighing the costs of storage versus resampling, the brain arguably favors storage when perceptual access requires an artificial action, but not when input is accessible through a natural exploratory mechanism such as eye movements.

This was also demonstrated by Li et al. (2023), who used a task similar to the present one, and compared a condition in which visual information was fully accessible to a condition in which it was revealed only when the computer mouse hovered above it. Consistent with our findings, they showed greater VWM use in the unnatural (mouse) condition than in the natural free-view condition. Although their findings align with our hypothesis, that study was not designed to isolate the effect of free exploration or to examine the role of eye movements in VWM utilization. First, eye movements were not restricted in the mouse condition and likely followed the cursor, resembling the free-view condition. Second, the free-view and mouse conditions differed not only in motor responses but also in the availability of visual information, which was fully visible in the free-view condition but largely occluded in the mouse condition. In our design, full access to visual information was provided in both conditions, but in one it was achieved through natural exploratory behavior (eye movements), and in the other through an artificial behavior (button press). This design allowed us to examine the link between the mean of accessing information and VWM utilization.

### 5.2. The effect of voluntary control on VWM utilization

Our finding that observers reached a VWM capacity of around 3-5 items in the sequential condition seemingly contrasts with the findings of Kvitelashvili & Kessler (2024), who used an adapted change detection task and showed that participants favored repeated perceptual sampling over reliance on VWM. In their study, switching was executed via button presses, as in our sequential condition, yet they observed underutilization of VWM. This difference can be explained by differences in task demands: our task required visual search and comparison between two equally rich displays, whereas theirs required recall and reconstruction. In their paradigm, switching often meant moving from a meaningful source of information (the model) to a blank or partially completed reconstruction screen. This reliance on precise recall and reproduction may have encouraged serial resampling rather than sustained memory storage. In contrast, our task allowed for partial encoding strategies - such as scanning for likely matches, which supported more flexible use of VWM depending on switching costs. This illustrates how task structure and stimulus can critically shape VWM strategies, underscoring the need for further research on the interaction between motor control and working memory.

### 5.3. The effect of items number on VWM Utilization

The present findings show that as the number of items increased, participants relied more on their VWM and less on perceptual resampling in both display conditions. This effect was stronger in the sequential than the simultaneous condition. The link between cognitive load and reliance on VWM is consistent with Draschkow et al., (2021), who used a visuomotor task in an immersive environment and found that reliance on VWM increased as task demands grew. Similarly, Droll et al. (2005) demonstrated that participants flexibly adapted their reliance on perception and VWM depending on task requirements: when a single feature was relevant, they relied primarily on perceptual input, but when multiple features were relevant, they stored information in advance. Together with the present results, these findings align with Baddeley’s (2007) model, which proposes that working memory engagement scales with cognitive demand. Our study extends this idea by showing that the relationship between load and VWM utilization depends on how observers access visual input, and that it is weaker with eye movements than button presses. This demonstrates the strong preference for resampling over storage in free viewing, even when cognitive load increases.

### 5.4. Visual search and foraging

The present findings are consistent with research on visual foraging tasks, in which participants search for multiple targets. Studies have examined the effect of eye movements on visual foraging by permitting or restricting eye movements in different groups while they were performing a visual search of multiple targets. Consistent with our findings, studies show that when observers can move their eyes freely, they exhibit more efficient oculomotor dynamics, which facilitate natural visual exploration and minimize memory load. In contrast, when eye movements are restricted, oculomotor behavior shifts toward a more ambient exploration mode, relying on covert attention and increased dependence on internal memory (Jóhannesson et al., 2016; Tagu & Kristjánsson, 2022). In the study by Jóhannesson et al. (2016), participants performed a novel multiple-target foraging task by either tapping targets with their fingers or fixating them with their eyes to compare how different response modalities affect attentional selection and foraging behavior. Results show notable differences between finger foraging and gaze foraging in how observers select multiple targets. Finger foraging showed a strong dissociation: participants switched easily between target categories when targets differed by a single feature but tended to make long runs of the same category during conjunction foraging, indicating more constrained switching under higher attentional demands. In contrast, gaze foraging demonstrated much less difference between feature and conjunction conditions, with a higher proportion of observers switching randomly between categories even under conjunction conditions. This reinforces our conclusion that eye movements are not merely the overt manifestation of attention, but they are the natural mechanism through which information is gathered and updated and, consequently, constraining them alters cognitive processes. Similar to our finding that free eye movements reduced reliance on VWM, foraging tasks demonstrate that overt exploration governs the balance between external sampling and internal maintenance. Our findings extend this literature by focusing on the utilization of working memory and showing that switching and gathering behaviors are modulated by eye movements, the natural means of information acquisition. Notably, our findings further reveal that eye movements can downregulate memory use even when perceptual input is fully available. Thus, while foraging paradigms highlight the behavioral coupling of gaze and selection, our findings reveal how this coupling modulates reliance on VWM.

### 5.5. Limitations and future directions

Our relatively small sample precluded analysis of inter-individual differences. Given robust variability in VWM capacity, future work should test how capacity relates to VWM use during free viewing. In addition, because we contrasted eye movements with button presses, we cannot speak to other exploratory actions, such as head or body movements. We hypothesize that these actions would show effects similar to eye movements, but this requires direct tests in future studies.

### 5.6. Conclusions

Findings of this study highlight the central role of visual exploration - one of the most natural and ubiquitous human behaviors - by showing that the ability to move the eyes freely shapes the balance between reliance on memory and perceptual input. This conclusion has far-reaching implications, given that much of cognitive research relies on paradigms that restrict eye movements. While fixation-based tasks provide valuable control, they may also distort our understanding of working memory and other cognitive processes (Hayhoe et al., 1998; Tatler & Land, 2011). Therefore, reconsidering some of the foundational assumptions of cognitive psychology in the context of naturalistic, free-viewing conditions is essential.

## Acknowledgement

This study was funded by ISF grant 2751/24 to S.Y-G, by BSF grant 2020308 to R.T.K. and S.Y-G. and Neubauer Family Foundation and Minducate scholarships to R.Q.

